# Ablation of MYB-dependent leukaemia phenotype in MLL-driven AML correlates with increased expression of MAFB

**DOI:** 10.1101/2020.05.27.118828

**Authors:** C Ward, P Cauchy, DS Walton, ML Clarke, D Blakemore, F Grebien, P Garcia, J Frampton, G Volpe

## Abstract

The transcription factor MYB plays a pivotal role in haematopoietic homeostasis and its aberrant expression is involved in the genesis and maintenance of acute myeloid leukaemia (AML). Our previous work has demonstrated that not all AML types display the same dependency on *MYB* expression and that MYB dependence is dictated by the nature of the driver mutation. However, whether this difference in MYB dependency is a general trend in AML still remains to be further elucidated. In this study, we investigate the importance of *MYB* in human leukaemia by performing siRNA-mediated knock-down in cell line models of AML with different driver lesions. We show that the characteristic reduction in proliferation and the concomitant induction of myeloid differentiation that is observed in MLL-fusion-driven leukaemia upon *MYB* suppression is not seen in AML cells with a complex karyotype. By performing transcriptome analysis, we demonstrate that a strong activation of *MAFB* expression driven by *MYB* ablation is restricted to MYB-dependent cells. In line with these observations, stratification of publicly available patient data reveals a reciprocal relationship between the expression of *MYB* and *MAFB*, highlighting a novel connection between those two factors in AML.

## INTRODUCTION

The MYB proto-oncoprotein is a sequence specific transcription factor that is highly expressed in immature haematopoietic progenitors and exerts master regulator functions during normal blood development. It controls the activation of genes that are necessary for definitive haematopoiesis, maintenance of stem cell self-renewal, and myeloid lineage specification and differentiation (Clarke et al., 2017; Lieu and Reddy, 2009; Mucenski et al., 1991; Sumner et al., 2000; Volpe et al., 2015).

Transcriptional dysregulation of MYB has been shown to be a central event in perpetuating a malignant self-renewal programme coupled with an arrest in normal myeloid differentiation, which are necessary events in the initiation and the progression of different types of leukaemia. The involvement of MYB in the development of myeloid diseases was originally demonstrated through the capacity of retrovirus-derived sequences to transform haematopoietic cells (Beug et al., 1979; Shen-Ong et al., 1986). This was followed by the identification of duplication of the MYB locus in paediatric acute lymphoblastic leukaemia (Clappier et al., 2007; Lahortiga et al., 2007) and of genomic rearrangements involving the MYB gene in acute basophilic and myelomonocytic leukaemia (Belloni et al., 2011; Murati et al., 2009; Quelen et al., 2011). Moreover, *MYB* has been shown to be an essential downstream target of multiple known oncogenic proteins such as HOXA9 and its TALE partners MEIS1 and PBX1 (Dasse et al., 2012; Hess et al., 2006). These latter factors are themselves downstream targets of MLL fusion proteins (Dasse et al., 2012; Hess et al., 2006) and *Myb* was demonstrated to be a strong mediator of oncogenic addiction in both cell lines and mouse models of MLL-AF9 driven leukaemia (Zuber et al., 2011). Previous studies from our group demonstrated that one mechanism through which MYB influences AML establishment involves the enforcement of sustained expression of *FLT3*, which is mediated through a functional cooperation with C/EBPα (Volpe et al., 2015; Volpe et al., 2013). Furthermore, we have also demonstrated that the dependency on *Myb* generally observed in leukaemia is attenuated in a murine model of AML that is driven by biallelic CEBPA N-terminal mutations (Volpe et al., 2019).

Here, we sought to assess in more detail how the leukaemia phenotype-related dependence on MYB translates to human leukaemia that are driven by different genetic lesions. In this pursuit, we focussed on human AML driven by either MLL-fusions versus those characterized by complex karyotypic lesions and assessed if and how those different leukaemia contexts would display a characteristic phenotypic modulation in response to *MYB* suppression. In agreement with previous reports, we show that depletion of *MYB* reverses the leukaemia-associated cellular phenotypes in MLL-fusion-driven AML, which is accompanied by de-repression of *MAFB*. However, we find that AML with complex karyotypic lesions does not undergo the conventional transcriptional and morphological alterations that are associated with MYB suppression. Gene expression data from cohorts of patients with either MLL-translocations versus complex karyotype were dichotomized based on the expression of either *MYB* or *MAFB*. In line with data obtained in cell lines, analysis of gene expression profiles showed that patients with low *MYB* expression and high *MAFB* levels display a similar expression signature.

## MATERIAL AND METHODS

### Cell lines

FUJIOKA and MOLM14 cells were cultured in RPMI-1640 medium supplemented with 10% foetal bovine serum (FBS), 50U/ml penicillin, 50µg/ml streptomycin, 2mM L-glutamine. Cell were maintained at 0.5 × 10^6^ cells/ml at 37°C with 5% CO_2_ in a humidified incubator and were washed with phosphate buffered saline solution between passages.

### Transfection experiments, proliferation and differentiation assays

In total, 5 × 10^6^ FUJIOKA or MOLM14 cells were electroporated using a EPI 3500 (Fischer, Germany) single 250 V pulse for 10ms with 300mM of *MYB* siRNA or a scrambled negative control siRNA (SIGMA ALDRICH) as previously reported in Clarke et al (Clarke et al., 2017). After electroporation, cells were kept in the electroporation cuvette for 10 minutes after which cells were added RPMI-1640 with 10% FBS, supplemented with penicillin/streptomycin and L-glutamine at a concentration of cells per ml and returned to an incubator kept at 37°C and 5% CO_2_. After transfection cells were plated at a density 10^6^ cells/ml and viable cells were counted and passaged at a ratio of 1:2 every 24 hours for four consecutive days to determine their proliferative capacity. Assessment of differentiation at 96 hours post *Myb* knockdown was achieved by flow cytometry staining of the cells with anti-CD11b PE-Cy7 (eBioscience). Acquisition and analysis of flow cytometric data was performed using Cyan ADP with Summit 4.4 software.

### Quantitative RT-PCR and Western Blot

10^6^ cells from both FUJIOKA and MOLM14 cells lines were collected 24h post transfection and RNA was extracted using RNeasy Mini kit (QIAGEN), and first-strand cDNA synthesis was performed using standard protocols. Quantitative RT-PCR analysis for *MYB* and *MAFB* was performed using predesigned Taqman gene expression assays (Applied Biosystems). Total protein lysates obtained from transfected cells were used for Western Blot analysis using the following antibodies: anti-MYB mouse monoclonal (1:1000, Upstate/Millipore) and anti-GAPDH mouse monoclonal (1:10000 dilution, Abcam).

### Statistical analysis

Statistical significance was performed by applying Student’s t-test for pairwise comparison and the p-values are indicated where appropriate. Analysis of *MYB* and *MAFB* expression in human patient microarray data from the Haferlach cohort presented in Figure 1A and 4A was performed using non-parametric Kruskal-Wallis test. All statistical analysis was performed using Graphpad Prism 7 (Graphpad Software Inc).

**Figure 1.**
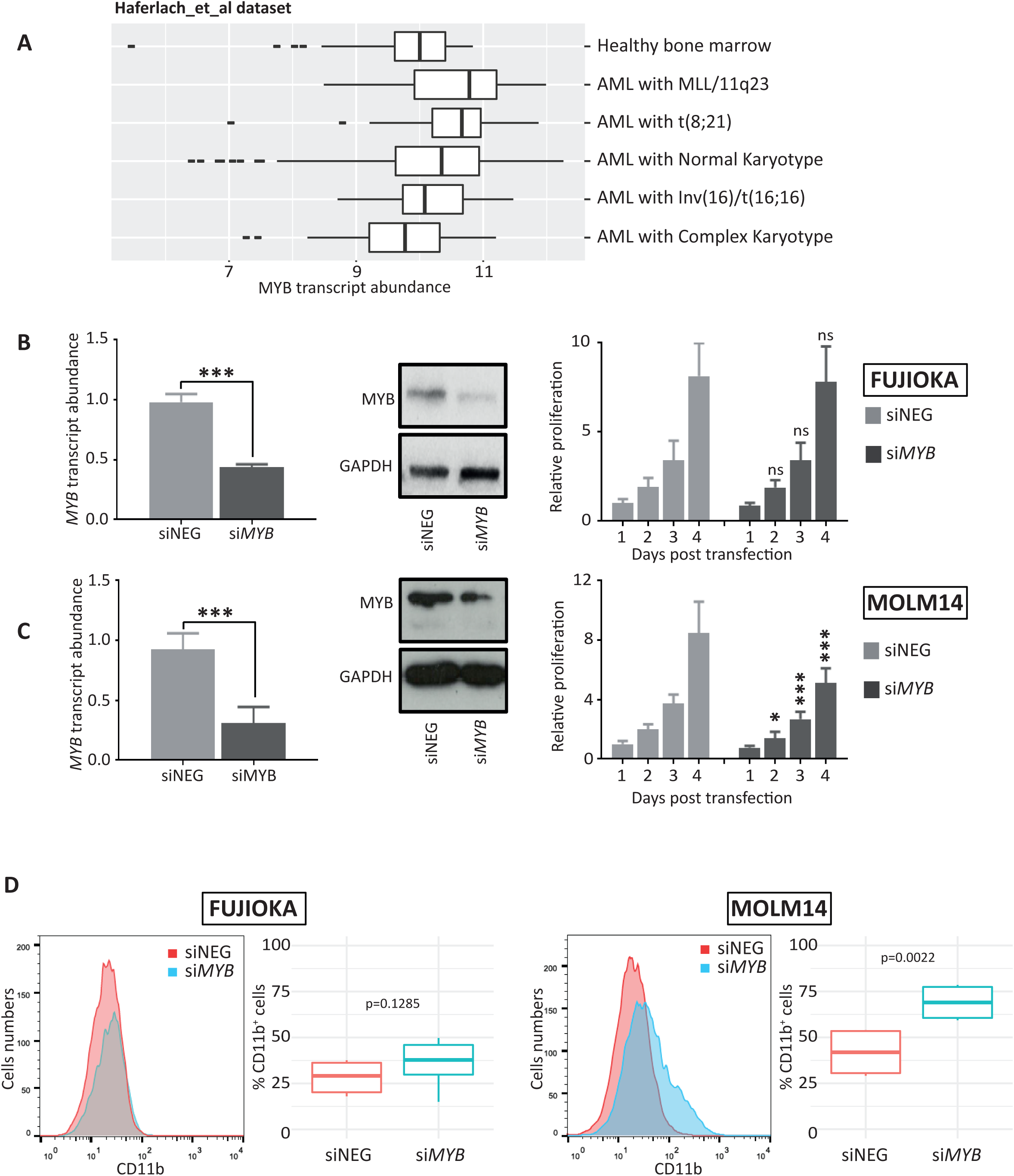
*MYB* depletion in AML driven by MLL-fusions reverses their leukaemia phenotype. **(A)** Boxplot depicting the abundance of *MYB* transcript in subgroups of patients from the dataset reported by Harferlach et al, characterized by the karyotypic abnormalities indicated in the graph in comparison with healthy bone marrow donors. The statistical significance presented in this plot has been determined by applying a Kruskal-Wallis t-test. **(B)** and **(C)** *MYB* knock-down efficiency in FUJIOKA and MOLM14 cells was determined 24h post transfection by quantitative-PCR and immunoblotting in cells transfected with *MYB* siRNA or scrambled negative control. Cell viability was determined by counting cells every 24h for 4 consecutive days. Results are representative of 3 independent experiments. Statistical analysis was calculated using Student’s t-test. (***, p<0.001 and **, p<0.01, *, p<0.05). **(D)** Histogram representing the analysis of CD11b myeloid surface marker expression in both cell lines transfected with either *MYB* siRNA or corresponding scrambled negative control. The boxplot represents the percentage of CD11b^+^ cells calculated from 3 independent experiments. The p-value indicated in the figure was calculated using Student’s t-test.

### RNA-sequencing

For RNA-Seq, libraries were prepared using the Illumina TruSeq Stranded kit according to the manufacturer’s instructions. Sequencing was performed in Genomics facility of the Institute of Cancer and Genomic Sciences, University of Birmingham, Birmingham, UK on an Illumina NextSeq 500 instrument.

### RNA-seq and differential gene expression analysis

Reads were trimmed using Trim Galore v 0.5.0 with --paired option. Gene counts were calculated using RSEM v1.2.22 using rsem-calculate-expression with --bowtie2, --bowtie2-sensitivity-level very_sensitive, --output-genome-bam and --paired-end options. The reads were aligned to hg38 assembly indexed with version 81 Ensembl GTF. Counts were normalized using DESeq2 v1.24 in R v3.6.0. Differential expression analysis was performed using the deseq function in DESeq2. Genes were considered significant if their adjusted p.value was less than 0.05 and their log2 fold change was greater than 2 or less than −2. Heatmaps were generated using the pheatmap v1.0.12 package and the pheatmap function. Euler diagrams were produced using the VennDiagram v1.6.20 package and the venn.diagram function using the intersected significantly up-regulated genes or the significantly down-regulated genes. Volcano plots were produced using the ggplot2 v3.2.1 package.

### Genome browser tracks

The genomic bam output from RSEM was used in the deeptools v3.1.3 bamCoverage function with --normalizeUsing CPM and the -bl hg38.blacklist.bed options. The resulting bigwig file was uploaded to a web server and visualised on UCSC genome browser.

### Patient profiling array and data processing

GSE13204 (Haferlach et al., 2010) raw data was downloaded from NCBI Gene Expression Omnibus (GEO). Patients were ranked for high or low *MYB* or *MAFB* expression using quantile cut off values as follows: 75^th^ percentile and higher were considered high expressers while 25^th^ percentile and lower were considered low expressers. Patients were subclassified and gene differential gene expression analysis was carried out as previously reported (Volpe et al., 2017; Ward et al., 2020).

### Data availability

RNA-Seq data generated in this study are available at the Gene Expression Omnibus (GEO) under series GSE149556.

## RESULTS

### Reduction of *MYB* expression does not reverse the block in complex karyotype AML cells

We have previously demonstrated that *Myb* is required for the maintenance of CEBPA-mutant AML and that the dependency on *Myb* expression is dictated by the nature of the mutations that drive the leukemic phenotype (Volpe et al., 2019; Volpe et al., 2013). To assess the relevance of our previous findings to human leukaemia and to test whether different leukaemia subtypes show a different dependency on *MYB* expression as observed in the case of CEBPA mutations, we analysed publicly available patient array data (Haferlach et al., 2010) by classifying patients based on their karyotypic abnormalities. In line with previous reports, we found *MYB* expression to be the highest in AML with MLL fusions and the t(8;21) translocation giving rise to the RUNX1-ETO oncoprotein when compared with healthy bone marrow donors. In contrast, *MYB* expression was not elevated in patients with Inv(16) lesions and in those with abnormal/complex karyotypes (Fig. 1A). To assess how reducing *MYB* levels would impact on the maintenance of leukaemia classes that are characterized by different levels of *MYB* expression, we chose to focus on leukaemia driven either by MLL fusions or complex karyotypes. We employed MOLM14, representing a well-characterized model of MLL-AF9 driven leukaemia, and FUJIOKA cells as a model for complex karyotype leukaemia.

To determine the immediate consequences of *MYB* reduction in those two classes of AML, we transfected the cell lines with siRNA targeting either *MYB* or a scrambled negative control. Efficient reduction in *MYB* expression (60-80%) was verified at both RNA (24h post transfection) and protein (48h post transfection) levels (Fig. 1B, 1C). Transfected cells were cultured for up to 96h, and cell numbers were determined daily to assess whether *MYB* reduction affects proliferation capacity. Decreased *MYB* levels led to a pronounced growth retardation in MOLM14, confirming the dependency of these cells on *MYB* expression (Zuber et al., 2011). Conversely, *MYB* suppression did not affect the proliferation rate of FUJIOKA cells (Fig. 1B, 1C). As we have previously demonstrated that *MYB* knockdown in different leukaemia phenotypes can override the differentiation block to instruct a myeloid commitment programme, we tested the differentiation capacity of cells by measuring CD11b surface marker expression by flow cytometry. While MOLM14 displayed a clear induction of myeloid differentiation, no upregulation of CD11b was seen in the complex karyotype cells (Fig. 1D).

Altogether, these results suggest that leukaemia with complex karyotype lacks a major dependency on *MYB* expression.

### Molecular consequences of *MYB* reduction in MLL-driven and complex karyotype AML cell lines

Previous studies have reported the capacity of *MYB* to influence leukaemia establishment and maintenance through enforcing a self-renewal programme while suppressing myeloid haematopoietic commitment (Lorenzo et al., 2011; Volpe et al., 2019; Zhao et al., 2011; Zuber et al., 2011). In order to understand how *MYB* reduction could lead to such a different responses in different leukaemia settings, we investigated the transcriptomes of MOLM14 and FUJIOKA cells 24h after *MYB* KD using RNA-sequencing. For this analysis, we considered up-regulated genes as those displaying an average Log2 fold change (FC) above 2 and the down-regulated ones with values below −2, with an adjusted p-value < 0.05. *MYB* knockdown did not induce any broad changes in gene expression in FUJIOKA cells, with only 2 genes being up-regulated and 2 genes being down-regulated. In contrast, we observed that siRNA-mediated knock-down of *MYB* in MOLM14 led to 13 down-regulated and 148 up-regulated genes, which is in agreement with previous reports of *MYB*-induced gene repression in myeloid cells (Fig. 2A). *MYB* knockdown in MOLM14 cells recapitulated the typical pattern of upregulation of myeloid genes that are known targets of *MYB*, such as *PIM1, ITGAM/CD11b, MYC, BCL2, DUSP6* and *GFI1* (Fig. 2B, S1A). Notably, this analysis also pointed at a large increase in the expression of *MAFB*, a transcription factor that has been previously associated with haematological malignancies (Fig. 2C) (Pajcini et al., 2017; Sieweke et al., 1997; Stralen et al., 2009). The striking up-regulation of *MAFB* in response to *MYB* reduction was also confirmed by quantitative PCR analysis in MOLM14 cells, while no significant difference was observed in FUJIOKA cells (Fig. 2D). Gene set enrichment analysis (GSEA) demonstrated that *MYB* reduction led to a down-regulation of the leukaemia stem cell (LSC) gene expression programme in MOLM14 cells but not in FUJIOKA cells (Fig. S1B).

**Figure 2.**
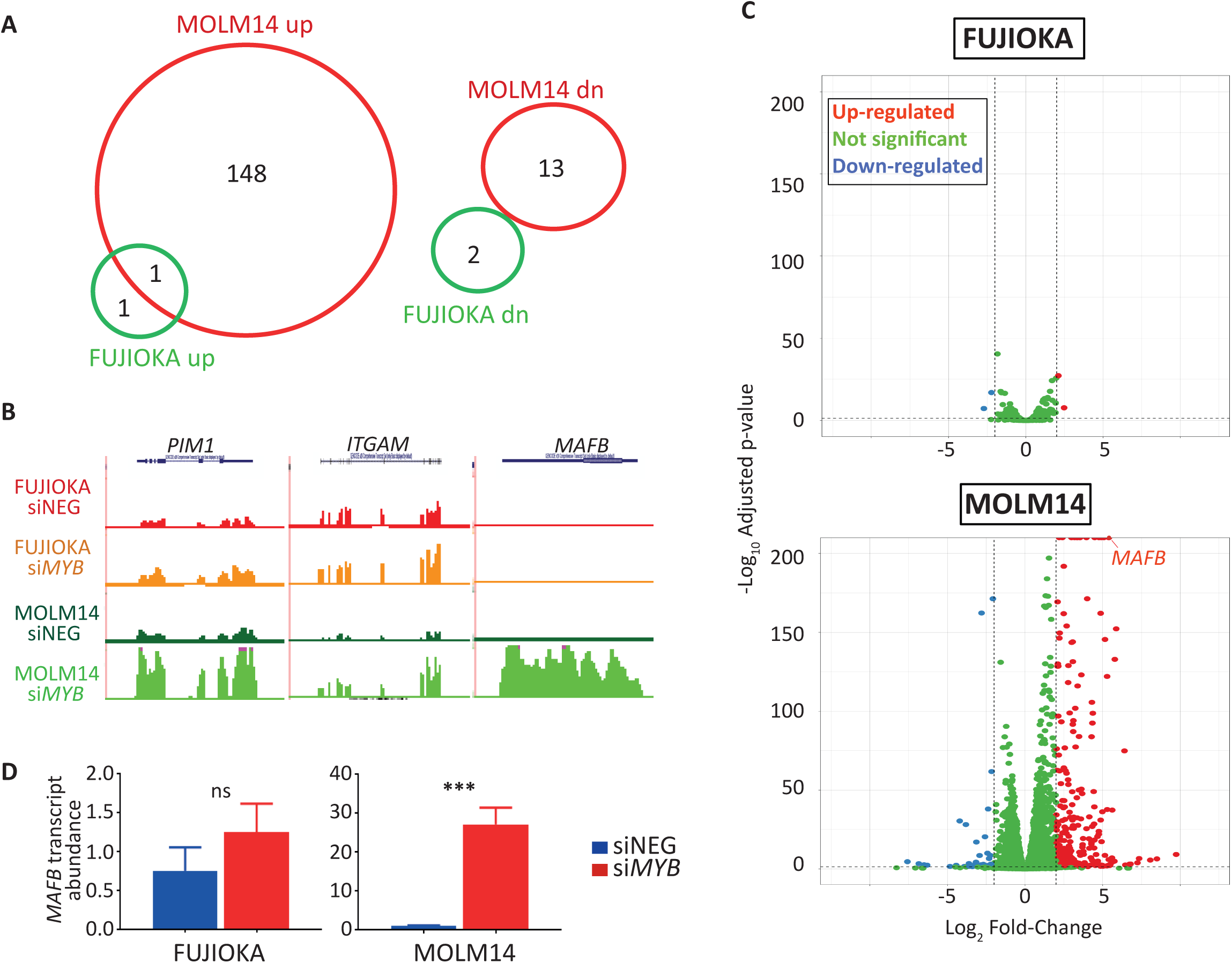
Molecular consequences of *MYB* depletion in human AML cell lines. **(A)** Euler diagrams indicating the numbers and overlap of genes up- and down-regulated in response to MYB suppression in both cell lines. **(B)** UCSC genome browser screenshots of RNA-seq performed in FUJIOKA and MOLM14 cells with either siNEG or siMYB treatment at *PIM1, ITGAM* and *MAFB* loci. Profiles are scaled to 1% GAPDH. **(C)** Volcano plot representing the differentially regulated genes in the response to MYB siRNA treatment. Up-regulated (over 2 log_2_ FC) and down-regulated (below −2 log_2_ FC) are indicated by red and blue dots, respectively. **(D)** qPCR analysis of *MAFB* transcript abundance in both cell lines transfected with either siNEG or si*MYB*. This analysis represents an average of 3 independent experiments. The statistical significance shown in the plot was calculated using Student’s t-test. (***, p<0.001; ns, not significant).

Our data suggests that one potential mechanism by which *MYB* reduction influences the phenotype in MLL-rearranged leukaemia could be through the activation of *MAFB* expression.

### Correlation of *MYB* and *MAFB* expression in human MLL-driven leukaemia

We next sought to confirm our findings obtained in cell lines by selecting subgroups of AML patients characterized by either abnormal karyotype or MLL fusions from a large cohort (Haferlach et al., 2010). For this purpose, we ranked patients in each subgroup according to *MYB* expression and selected the bottom and the top quartiles of the whole expression range as low and high expressers, respectively (Fig. 3A). We performed gene expression analysis to identify similarities and differences between the subgroups by focussing on differentially expressed mRNAs with a log2 FC below −2 or above 2; by doing so, we identified 204 versus 111 genes that displayed a positive correlation with *MYB* expression and 217 versus 14 genes that displayed a negative correlation in the subgroups characterized by MLL aberrations and complex karyotype, respectively (Fig. 3B). These findings are in agreement with our experimental observations, as the transcriptomic differences observed when comparing *MYB* low vs high MLL patients show remarkably high correlation with the transcriptomic perturbation seen in MOLM14 upon *MYB* suppression. Conversely, only minor differences were observed in complex karyotype patients, thus mirroring our FUJIOKA experimental findings. Importantly, we observed *MAFB* to be among the most negatively correlated genes (−5.7 FC). Intersection of transcriptome differences within the subgroups of patients showed only a minimal overlap, this being 1.38% versus 21% and 12% versus 22% for the negatively and positively correlated genes, respectively (Fig. 3C), which is consistent with the observation that MYB reduction in different AML subgroups can lead to different transcriptional outcomes. To investigate this in more detail, we screened the subgroups of patients by looking at the expression of typical myeloid genes that are known MYB targets; we found marked differences in the expression of several genes that are normally either repressed (e.g. *BCL2, CDK6, FLT3, GFI1 and KIT*) or activated (such as *BCL6, CD14, DUSP6 and ITGAM*) by MYB, in MLL-rearranged leukaemia patients, while only minimal or no difference was observed for the same genes in the complex karyotype subgroup when comparing MYB-high versus MYB-low subgroups (Fig. 3D).

**Figure 3.**
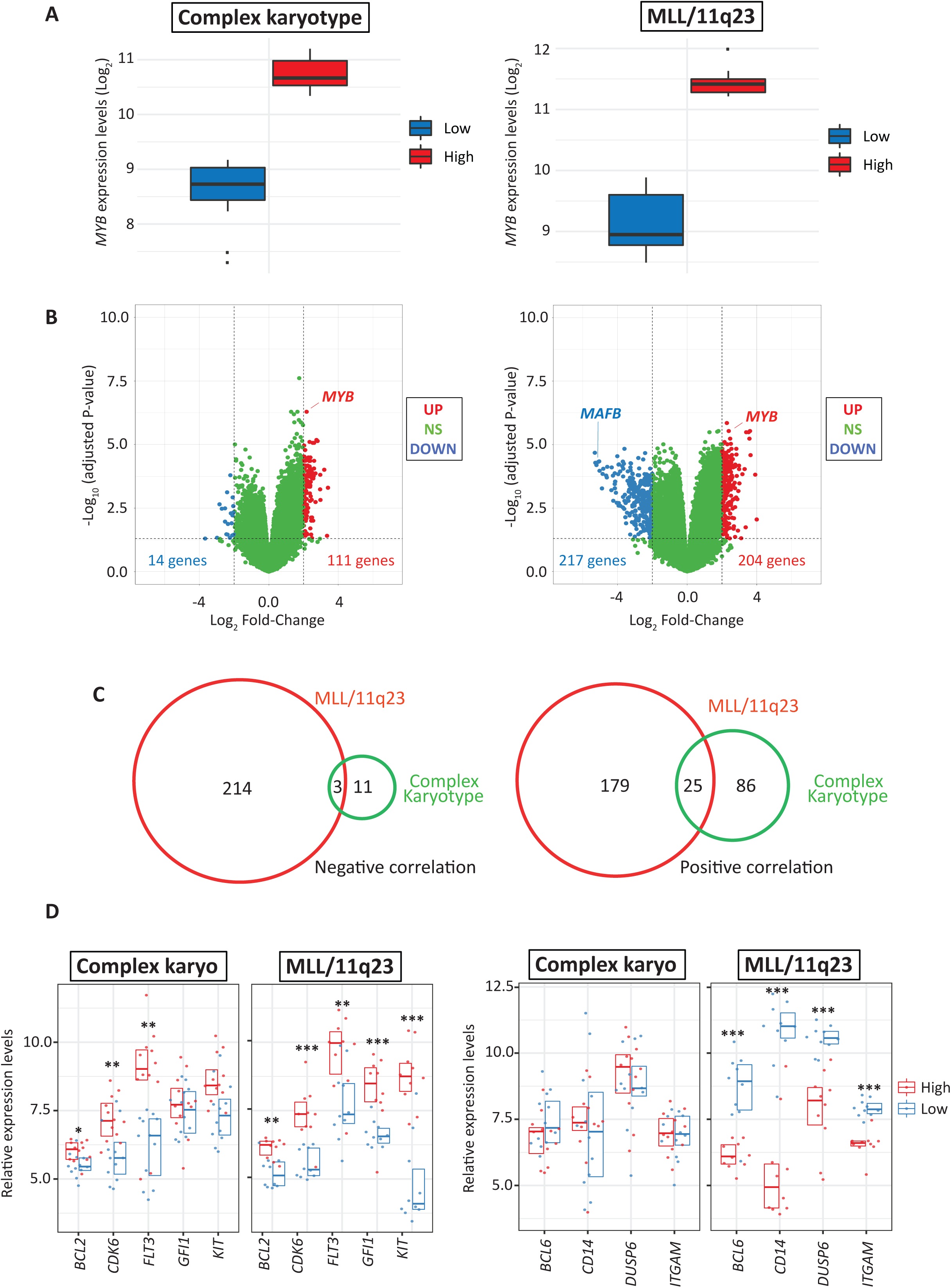
Validation of siRNA-mediated *MYB* knock-down consequences in AML patient expression arrays. **(A)** Boxplot representing the boundaries of *MYB*^low^ (lower quartile, 0-25% of expression range) and *MYB*^high^ (upper quartile, 75-100% of expression range) patients in the subgroups with complex karyotype (left panel) or MLL-translocations (right panels) from the cohort reported by Haferlach et al. **(B)** Volcano plot representing differentially expressed genes that display either negative (below −2 log_2_ FC, indicated as blue dots) or positive (above 2 log_2_ FC, indicated as red dots) correlation when comparing high versus low expressers in each AML subgroup. **(C)** Euler diagrams showing the numbers and overlap of genes that are either negatively (left panel) or positively (right panel) correlated with *MYB* expression when comparing complex karyotype and MLL-fusion subgroups. **(D)** Expression of representative MYB myeloid target genes separated in positive (*BCL2, CDK6, FLT3, GFI1* and *KIT*) vs negative regulated genes (*BCL6, CD14, DUSP6* and *ITGAM*) in both subgroups. Data are presented as overlapping scatter plots in which *MYB*^high^ patients are indicated in red and *MYB*^low^ patients are indicated in blue. Every plot shows a color-coded boxplot showing a median interquartile range. The statistical analysis shown in each plot indicates p-value adjusted for false discovery rates (***<0.001, **<0.01, *<0.05).

Our correlative analysis further reinforces the idea that different classes of leukaemias display different dependencies of MYB expression.

### High *MAFB* expression phenocopies the transcriptional consequences of lower *MYB* expression in human AML patients with MLL rearrangements

To further validate the reciprocal association between *MAFB* and *MYB* expression, we screened the subgroups of patients from the Haferlach cohort for the expression of *MAFB*. This analysis pointed out that patients carrying an MLL translocation, which are characterized by the highest *MYB* expression within the whole cohort, are among those with the lowest *MAFB* RNA abundance (Fig. 4A).

**Figure 4.**
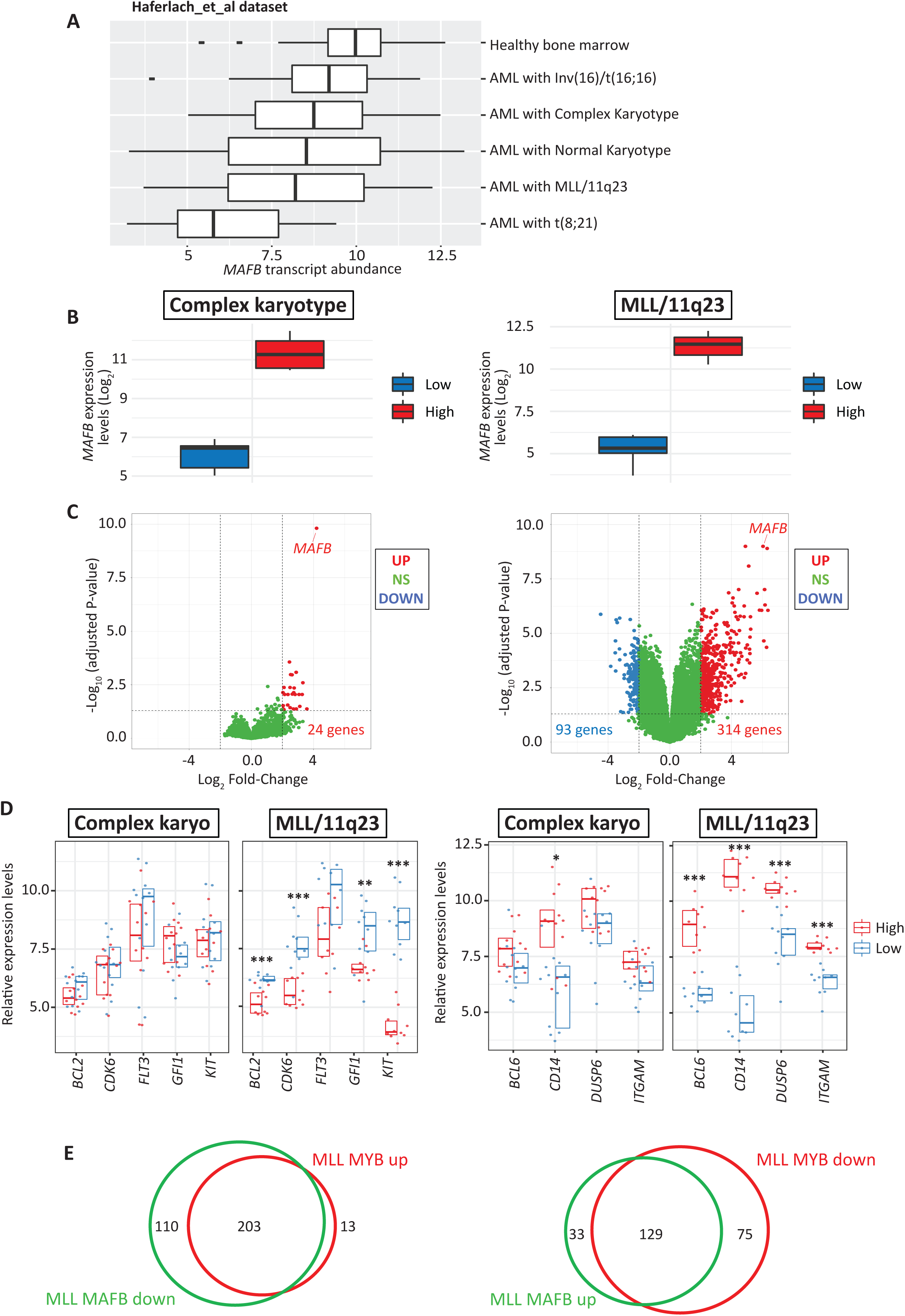
Correlation of the molecular consequences of high *MAFB* and low *MYB* in AML patients with MLL translocations. **(A)** Boxplot depicting the abundance of *MAFB* mRNA in subgroups of patients from the Haferlach dataset, characterized by the karyotypic abnormalities indicated in the graph in comparison with healthy bone marrow donors. The statistical significance presented in this plot has been determined by applying a Kruskal-Wallis t-test. **(B)** Boxplot representing the boundaries of *MAFB*^low^ (lower quartile, 0-25% of expression range) and *MAFB*^high^ (upper quantile, 75-100% of expression range) patients in the subgroups with complex karyotype (left panel) or MLL-translocations (right panels) from the Haferlach cohort. **(C)** Volcano plot indicating differentially expressed mRNAs with either negative (below −2 log_2_ FC, indicated as blue dots) or positive (above 2 log_2_ FC, indicated as red dots) correlation when comparing high versus low *MAFB* expressers in each AML subgroup. **(D)** MYB myeloid target genes separated in positive (*BCL2, CDK6, FLT3, GFI1* and *KIT*) vs negative regulated genes (*BCL6, CD14, DUSP6* and *ITGAM*) comparing low versus high expressers in both complex karyotype and MLL-translocation subgroups. Data are presented as overlapping scatter plots in which *MAFB*^high^ patients are indicated in red and *MAFB*^low^ patients are indicated in blue. Every plot shows a color-coded boxplot showing a median interquartile range. The statistical analysis shown in each plot indicates p-value adjusted for false discovery rates (***<0.001, **<0.01, *<0.05). **(E)** Euler diagrams indicating the numbers and overlap of genes that display negative correlation with *MAFB* and positive correlation with *MYB* (left panel) or genes that display positive correlation with *MAFB* and negative correlation with *MYB* (right panel) in the subgroup of patients carrying MLL translocations.

In order to compare the consequences of different *MAFB* levels, we ranked patients from the MLL and complex karyotype subgroups according to *MAFB* expression and selected the bottom and the top quartile of the whole range as low and high expressers, respectively (Fig. 4B). We next performed differential gene expression analysis using the same parameters described for the *MYB* classification and observed a large number of differentially regulated transcripts in the patients of the subset with MLL fusions (314 positively and 93 negatively correlated genes) while only a small set of genes was differentially regulated in AML patients with a complex karyotype (24 positively and no negatively correlated genes) (Fig. 4C).

Having demonstrated a significant correlation between *MYB* and *MAFB* levels, we next investigated the relationship between high *MAFB* expression and known target genes of MYB in the context of myeloid cells (as described in Fig. 3D). In patients with MLL rearrangements, genes that are positively correlated with *MYB* are generally negatively correlated with *MAFB*, expression, such as *BCL6, CD14, DUSP6* and *ITGAM*. Conversely, genes that are negatively correlated with *MYB* showed a positive correlation with *MAFB* expression, those being *BCL2, CDK6, FLT3, GFI1* and *KIT* (Fig. 4D). Furthermore, we observed a generally large overlap between the genes whose expression is positively correlated with *MYB* and negatively correlated with *MAFB* and between genes that are negatively correlated with by MYB and positively correlated with MAFB (Fig. 4E).

In summary, our data highlight a reciprocal correlation between the expression of *MYB* and *MAFB* in MLL-fusion-driven AML, suggesting that *MAFB* could be a new read-out for the response of *MYB* depletion or inhibition in this disease subtype. In addition, these comparisons further demonstrate that the way in which MYB influences the leukaemia phenotype depends on the driving mutation.

## DISCUSSION

In the present study, we used siRNA-mediated *MYB* silencing and transcriptome profiling to explore the oncogenic addiction to *MYB* levels of human leukaemia driven by different genetic lesions. By taking this approach in cell lines modelling the main leukaemia classes, that is MLL-fusions and complex karyotypes, we have shown that while MLL-fusion-driven AML requires MYB to enforce self-renewal and a myeloid differentiation barrier, complex karyotype leukaemia shows little or no dependency on *MYB* expression.

In line with previous observations from our work on C/EBPα mutant models, we find that the phenotypic response to *MYB* depletion is very different in the MLL-fusion-driven MOLM-14 cell line as compared to FUJIOKA cells, which represent a model for complex karyotype AML, and that this is reflected in distinct modulations of the transcriptome. In fact, we show that perturbation of *MYB* expression is able to reverse the differentiation block normally observed in leukaemia and impairs the self-renewal capacity of MOLM14 cells, while such a reduction in *MYB* appears to be well tolerated in FUJIOKA cells as the undifferentiated state persists and the differentiation block is not overcome.

The extensive changes in gene expression seen in MLL-fusion-driven AML cells in response to *MYB* depletion include a decrease in the leukaemia stem cell signature and modulation of many genes that are known to be targets of MYB, including *PIM1, ITGAM, GFI1, FLT3, CD14*, and *DUSP6*. Most interestingly, MYB depletion is accompanied by a striking increase in the abundance of *MAFB* RNA. A role for MafB in the normal promotion of myeloid differentiation is well established (Kelly et al., 2000) and it is known that Myb can inhibit MafB transactivation potential through direct binding to a SUMOylated form of MafB (Tillmanns et al., 2007). Taken together, this suggests that MYB normally supports the leukaemia phenotype in MLL-rearranged AML by directly or indirectly limiting *MAFB* expression, with the additional possibility of suppression of MAFB protein activity, thereby limiting the ability of the cells to differentiate. In our study, we also sought to determine whether elevation of *MAFB* levels would phenocopy the consequences of MYB inhibition; as such, we performed ectopic expression though lentiviral infection in FUJIOKA and MOLM14 cells, though we only achieved disappointing results as we failed to obtain sufficient infection rates to observe an effect on those cells.

The lack of dependency of the leukaemia phenotype on *MYB* levels in complex karyotype AML cells, represented in our study by FUJIOKA cells, is mirrored by a small number of changes in gene expression upon *MYB* depletion in this cell line. Strikingly, no changes in the levels of *MAFB* or known MYB targets such as *GFI1, FLT3*, and *PIM1* were observed. The fact that these latter genes are known for their capacity to enforce self-renewal and a myeloid commitment block provides a hint as to why complex karyotype AML might show very minimal response to *MYB* depletion.

To strengthen the validity of our observations, we performed bioinformatic analysis of publicly available AML patient gene expression datasets. We stratified patients characterized by either complex karyotype or MLL-fusions based on *MYB* or *MAFB* mRNA levels. By comparing low versus high *MYB* expressers, we found that the transcriptome differences paralleled our findings obtained through experimental manipulation of *MYB* in the MOLM14 cells line, with a large number of genes positively and negatively correlated with MYB levels, including well-known MYB targets that are normally activated (*BCL2, CDK6, FLT3 and GFI1*) or repressed (*BCL6, CD14, ITGAM and DUSP6*) in different cellular contexts. Importantly, this analysis highlighted *MAFB* as a gene whose expression was the most negatively correlated with *MYB* RNA levels, thus further highlighting a significant connection between these two genes. In contrast to the strong *MYB* level-associated gene expression differences seen in MLL-driven AML, only a small number of differentially expressed genes was found in patients characterized by complex karyotypes. Also, this is in agreement with our observations in the FUJIOKA cells line, which exhibited only 2 down-regulated and 2 up-regulated genes in response to *MYB* depletion. This further underlines the idea that complex karyotype leukaemia has reduced or no dependency of *MYB* expression.

Given that *MAFB* levels appeared to be strongly associated with *MYB* expression, we also performed gene expression analysis of patient data by stratifying those samples based on the levels of *MAFB* RNA. Interestingly, this produced a large number of both positively and negative correlated genes in MLL-rearranged AML, while very few genes were found to be dependent upon *MAFB* expression in complex karyotype patients. Importantly, this analysis showed that genes whose expression is negatively correlated with *MYB* were strongly overlapping with genes whose expression is positively correlated with the level of *MAFB* RNA. Conversely, genes positively correlated with *MYB* expression were largely overlapping with those that are negatively correlated with *MAFB*. This was also confirmed by the significant differences detected for known *MYB* target genes.

In conclusion, this study sheds further light on the dependency of different categories of AML on the expression levels of the MYB transcription factor. We demonstrate that, as we had previously described in the case of models of leukaemia driven by mutations in C/EBPα, there can be vastly different responses to reduction in *MYB* levels that depend on the nature of the driving mutation. The finding of a strong inverse correlation between *MYB* and *MAFB* in MLL-driven leukaemia suggests that *MAFB* could serve both as a biomarker for prognosis and responsiveness to novel therapeutic strategies aimed at targeting MYB.

## AUTHOR CONTRIBUTIONS

GV conceived and designed the research and performed most of the experiments; CW and PC performed the bioinformatic analysis; DSW and MLC conducted the manipulation experiments; DB performed the immunoblotting. JF, MAEB, FG and PG provided relevant advice and financial support and edited the manuscript; GV and JF supervised the research. GV wrote and edited the manuscript.

## ACKNOWLEDGMENTS

CW is supported by Chinese Academy of Sciences President’s International Fellowship Initiative for Postdoctoral Researchers (2019PB0177) and a National Natural Science Foundation of China (31900466, 31900582), Natural Science Foundation of Guangdong Province, China (2018A030313379) and Research Fund for International Young Scientists grant (31950410553). GV is supported by Chinese Academy of Sciences President’s International Fellowship for Foreign Experts (2020FSB0002). This work was supported through funding provided by the College of Medical and Dental Sciences of the University of Birmingham and the Birmingham CRUK Centre.

---

**Figure S1.**
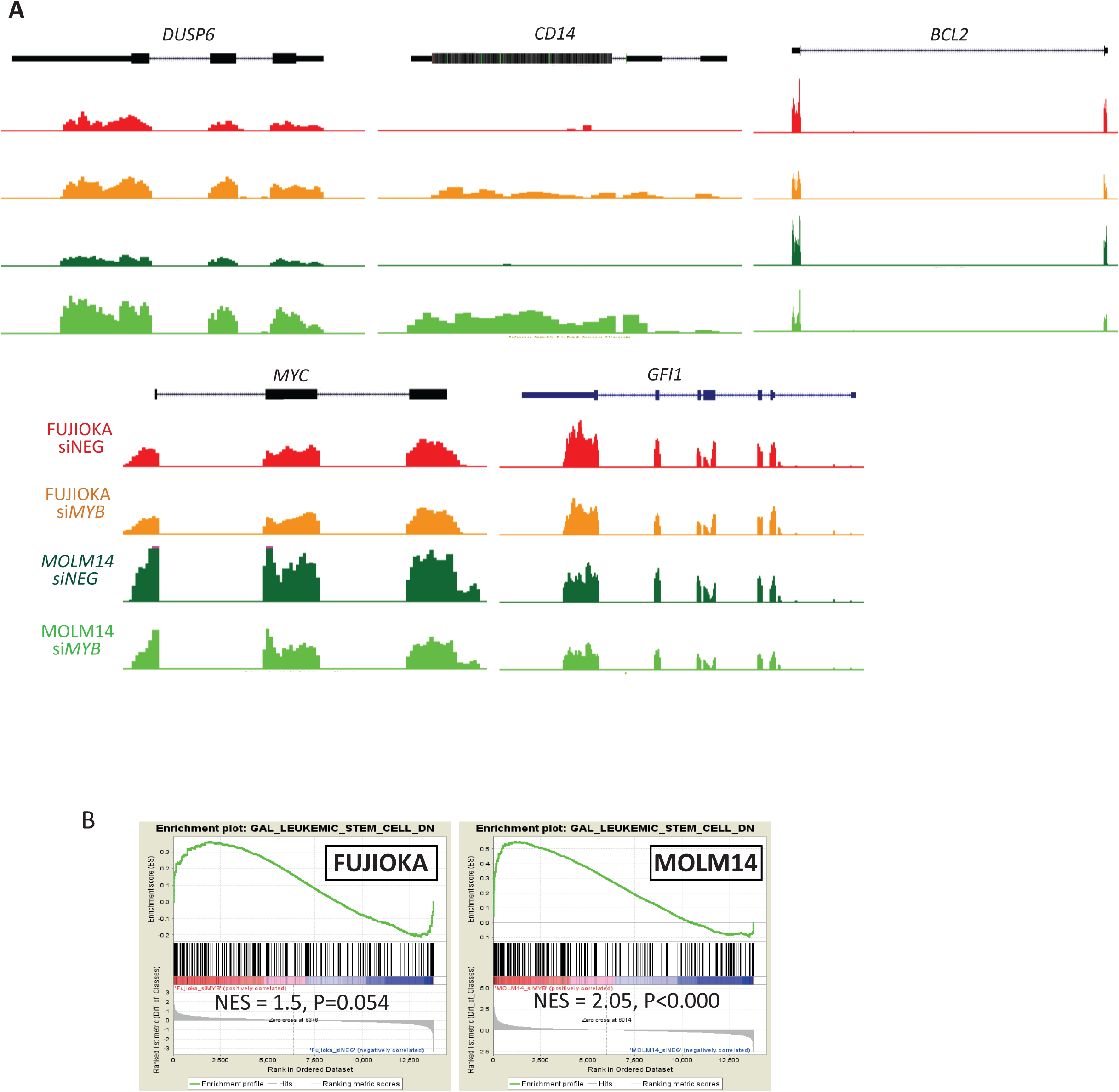
Gene expression consequences of *MYB* depletion. **(A)** UCSC genome browser screenshots of RNA-seq performed in FUJIOKA and MOLM14 cells with either siNEG or si*MYB* treatment at *DUSP6, CD14, BCL2, MYC*, and *MAFB* loci. Profiles are scaled to 1% GAPDH. (B) Gene set enrichment analysis of leukaemia stem cell signature genes in FUJIOKA and MOLM14 cells.

## REFERENCES

Belloni, E., Shing, D., Tapinassi, C., Viale, A., Mancuso, P., Malazzi, O., Gerbino, E., Dall’Olio, V., Egurbide, I., Odero, M.D., et al. (2011). In vivo expression of an aberrant MYB-GATA1 fusion induces leukemia in the presence of GATA1 reduced levels. Leukemia 25, 733–736.

Beug, H., von Kirchbach, A., Doderlein, G., Conscience, J.F., and Graf, T. (1979). Chicken hematopoietic cells transformed by seven strains of defective avian leukemia viruses display three distinct phenotypes of differentiation. Cell 18, 375–390.

Clappier, E., Cuccuini, W., Kalota, A., Crinquette, A., Cayuela, J.M., Dik, W.A., Langerak, A.W., Montpellier, B., Nadel, B., Walrafen, P., et al. (2007). The C-MYB locus is involved in chromosomal translocation and genomic duplications in human T-cell acute leukemia (T-ALL), the translocation defining a new T-ALL subtype in very young children. Blood 110, 1251–1261.

Clarke, M., Volpe, G., Sheriff, L., Walton, D., Ward, C., Wei, W., Dumon, S., Garcia, P., and Frampton, J. (2017). Transcriptional regulation of SPROUTY2 by MYB influences myeloid cell proliferation and stem cell properties by enhancing responsiveness to IL-3. Leukemia 31, 957–966.

Dasse, E., Volpe, G., Walton, D.S., Wilson, N., Del Pozzo, W., O’Neill, L.P., Slany, R.K., Frampton, J., and Dumon, S. (2012). Distinct regulation of c-myb gene expression by HoxA9, Meis1 and Pbx proteins in normal hematopoietic progenitors and transformed myeloid cells. Blood Cancer J 2, e76.

Haferlach, T., Kohlmann, A., Wieczorek, L., Basso, G., Kronnie, G.T., Bene, M.C., De Vos, J., Hernandez, J.M., Hofmann, W.K., Mills, K.I., et al. (2010). Clinical utility of microarray-based gene expression profiling in the diagnosis and subclassification of leukemia: report from the International Microarray Innovations in Leukemia Study Group. J Clin Oncol 28, 2529–2537.

Hess, J.L., Bittner, C.B., Zeisig, D.T., Bach, C., Fuchs, U., Borkhardt, A., Frampton, J., and Slany, R.K. (2006). c-Myb is an essential downstream target for homeobox-mediated transformation of hematopoietic cells. Blood 108, 297–304.

Kelly, L.M., Englmeier, U., Lafon, I., Sieweke, M.H., and Graf, T. (2000). MafB is an inducer of monocytic differentiation. EMBO J 19, 1987–1997.

Lahortiga, I., De Keersmaecker, K., Van Vlierberghe, P., Graux, C., Cauwelier, B., Lambert, F., Mentens, N., Beverloo, H.B., Pieters, R., Speleman, F., et al. (2007). Duplication of the MYB oncogene in T cell acute lymphoblastic leukemia. Nat Genet 39, 593–595.

Lieu, Y.K., and Reddy, E.P. (2009). Conditional c-myb knockout in adult hematopoietic stem cells leads to loss of self-renewal due to impaired proliferation and accelerated differentiation. Proc Natl Acad Sci U S A 106, 21689–21694.

Lorenzo, P.I., Brendeford, E.M., Gilfillan, S., Gavrilov, A.A., Leedsak, M., Razin, S.V., Eskeland, R., Saether, T., and Gabrielsen, O.S. (2011). Identification of c-Myb Target Genes in K562 Cells Reveals a Role for c-Myb as a Master Regulator. Genes Cancer 2, 805–817.

Mucenski, M.L., McLain, K., Kier, A.B., Swerdlow, S.H., Schreiner, C.M., Miller, T.A., Pietryga, D.W., Scott, W.J., Jr., and Potter, S.S. (1991). A functional c-myb gene is required for normal murine fetal hepatic hematopoiesis. Cell 65, 677–689.

Murati, A., Gervais, C., Carbuccia, N., Finetti, P., Cervera, N., Adelaide, J., Struski, S., Lippert, E., Mugneret, F., Tigaud, I., et al. (2009). Genome profiling of acute myelomonocytic leukemia: alteration of the MYB locus in MYST3-linked cases. Leukemia 23, 85–94.

Pajcini, K.V., Xu, L., Shao, L., Petrovic, J., Palasiewicz, K., Ohtani, Y., Bailis, W., Lee, C., Wertheim, G.B., Mani, R., et al. (2017). MAFB enhances oncogenic Notch signaling in T cell acute lymphoblastic leukemia. Sci Signal 10.

Quelen, C., Lippert, E., Struski, S., Demur, C., Soler, G., Prade, N., Delabesse, E., Broccardo, C., Dastugue, N., Mahon, F.X., et al. (2011). Identification of a transforming MYB-GATA1 fusion gene in acute basophilic leukemia: a new entity in male infants. Blood 117, 5719–5722.

Shen-Ong, G.L., Morse, H.C., 3rd, Potter, M., and Mushinski, J.F. (1986). Two modes of c-myb activation in virus-induced mouse myeloid tumors. Mol Cell Biol 6, 380–392.

Sieweke, M.H., Tekotte, H., Frampton, J., and Graf, T. (1997). MafB represses erythroid genes and differentiation through direct interaction with c-Ets-1. Leukemia 11 Suppl 3, 486–488.

Stralen, E., Leguit, R.J., Begthel, H., Michaux, L., Buijs, A., Lemmens, H., Scheiff, J.M., Doyen, C., Pierre, P., Forget, F., et al. (2009). MafB oncoprotein detected by immunohistochemistry as a highly sensitive and specific marker for the prognostic unfavorable t(14;20) (q32;q12) in multiple myeloma patients. Leukemia 23, 801–803.

Sumner, R., Crawford, A., Mucenski, M., and Frampton, J. (2000). Initiation of adult myelopoiesis can occur in the absence of c-Myb whereas subsequent development is strictly dependent on the transcription factor. Oncogene 19, 3335–3342.

Tillmanns, S., Otto, C., Jaffray, E., Du Roure, C., Bakri, Y., Vanhille, L., Sarrazin, S., Hay, R.T., and Sieweke, M.H. (2007). SUMO modification regulates MafB-driven macrophage differentiation by enabling Myb-dependent transcriptional repression. Mol Cell Biol 27, 5554–5564.

Volpe, G., Cauchy, P., Walton, D.S., Ward, C., Blakemore, D., Bayley, R., Clarke, M.L., Schmidt, L., Nerlov, C., Garcia, P., et al. (2019). Dependence on Myb expression is attenuated in myeloid leukaemia with N-terminal CEBPA mutations. Life Sci Alliance 2.

Volpe, G., Clarke, M., Garcia, P., Walton, D.S., Vegiopoulos, A., Del Pozzo, W., O’Neill, L.P., Frampton, J., and Dumon, S. (2015). Regulation of the Flt3 Gene in Haematopoietic Stem and Early Progenitor Cells. PLoS One 10, e0138257.

Volpe, G., Walton, D.S., Del Pozzo, W., Garcia, P., Dasse, E., O’Neill, L.P., Griffiths, M., Frampton, J., and Dumon, S. (2013). C/EBPalpha and MYB regulate FLT3 expression in AML. Leukemia 27, 1487–1496.

Volpe, G., Walton, D.S., Grainger, D.E., Ward, C., Cauchy, P., Blakemore, D., Coleman, D.J.L., Cockerill, P.N., Garcia, P., and Frampton, J. (2017). Prognostic significance of high GFI1 expression in AML of normal karyotype and its association with a FLT3-ITD signature. Sci Rep 7, 11148.

Ward, C., Cauchy, P., Garcia, P., Frampton, J., Esteban, M.A., and Volpe, G. (2020). High WBP5 expression correlates with elevation of HOX genes levels and is associated with inferior survival in patients with acute myeloid leukaemia. Scientific reports 10, 3505.

Zhao, L., Glazov, E.A., Pattabiraman, D.R., Al-Owaidi, F., Zhang, P., Brown, M.A., Leo, P.J., and Gonda, T.J. (2011). Integrated genome-wide chromatin occupancy and expression analyses identify key myeloid pro-differentiation transcription factors repressed by Myb. Nucleic Acids Res 39, 4664–4679.

Zuber, J., Rappaport, A.R., Luo, W., Wang, E., Chen, C., Vaseva, A.V., Shi, J., Weissmueller, S., Fellmann, C., Taylor, M.J., et al. (2011). An integrated approach to dissecting oncogene addiction implicates a Myb-coordinated self-renewal program as essential for leukemia maintenance. Genes Dev 25, 1628–1640.

